# A knowledge integration strategy for the selection of a robust multi-stress biomarkers panel for *Bacillus subtilis*

**DOI:** 10.1101/2022.06.20.496894

**Authors:** Yiming Huang, Nishant Sinha, Anil Wipat, Jaume Bacardit

## Abstract

Recent advances in high-throughput omics technologies have enhanced the identification of molecular biomarkers specific to phenotypes or states in bacteria. Using these biomarkers to monitor the state of bacteria used in biotechnological processes promises to increase process efficiency. However, live-cell monitoring systems applied to recognise bacterial cellular states in real time can only accommodate a small number of gene expression biomarkers. Computational methods are required to identify and prioritise robust biomarkers for experimental characterisation and verification. This study focused on designing a knowledge integration strategy for the selection of an optimal minimised gene expression biomarker panel to sense various stress states in *Bacillus subtilis*. We developed a computational method that ranks the candidate biomarker panels based on complementary information from machine learning model, gene regulatory network and co-expression network. We identified a recommended biomarker panel showing high stress sensing power for a variety of conditions included in both the dataset used for biomarker identification (mean f1-score achieved at 0.99) and the independent datasets from different sources (mean f1-score achieved at 0.98). We discovered a significant correlation between stress sensing power and evaluation metrics such as the number of associated regulators in a *B. subtilis* gene regulatory network (GRN) and the number of associated modules in a *B. subtilis* co-expression network (CEN). GRNs and CENs provide information relevant to the diversity of biological processes encoded by biomarker genes. We demonstrate that quantitatively relating meaningful evaluation metrics with stress sensing power has potential for recognising biomarkers that show better sensitivity and robustness to an extended set of stress conditions. We conclude that this approach is readily applicable to biomarker discovery model selection.

## 1 Introduction

Bioengineering applications often use bacteria as the host organisms to create high-value products such as pharmaceuticals, biofuels, fine chemicals, etc. [1]. While grown under optimal user-defined conditions, bacteria are still subjected to periods of cellular stresses that may lead to suboptimal growth and lower yields of target products [2-4] One of the goals of the biotechnology industry is to recognise these detrimental cellular states so that strategies for mitigating the damage can be engineered [5, 6]. Detrimental states can be characterised and recognised using a variety of techniques, from morphological and biochemical assays [7, 8] to omics methods [9]. In the case of the latter, panels of genes whose expression are indicative of certain cellular states can be used as transcriptional biomarkers.

By measuring the expression of a few key biomarker genes using amplicon panels[10, 11] or qPCR [12, 13], instead of characterising the global transcription using genome-scale RNA-seq or Microarray, the cellular states can be assayed with reduced costs. Moreover, while these sequencing measures are not easily carried out in real-time, having cellular state biomarkers is particularly attractive for single-cell measurement technologies that seek to develop ‘live cell’ biosensors using flow cytometry [14, 15] or microfluidics systems [16, 17]. These ‘live cell’ biosensors require the biomarker panels to be small, consisting of only a few key genes. This is because live monitoring of cellular state currently relies on the use of reporter genes, for which there are a limited number of distinct systems available.

Transcriptomic methods such as microarrays and RNA-seq are used to reveal gene expression patterns under different environmental conditions. These experiments can provide the genome-scale data necessary to identify transcriptional biomarkers for distinguishing physiological and biochemical states in a bacterial cell. By running statistical tests such as differential expression analysis, researchers have discovered the biomarker genes indicative of certain stress states for model bacteria used in the biotechnology industry, e.g., *Escherichia coli* [18, 19] and *Bacillus subtilis* [20, 21]. More biomarker studies in biomedical domains [22-24] have exploited the massive transcriptomics data by applying machine learning methods such as classification and feature selection models [25]. We recently proposed a pipeline of machine learning models to extract diverse cellular states from condition-dependent transcriptomics data and identify gene biomarkers that can classify different stress states for *B. subtilis* [26].

As transcriptomics data are characterised by high dimensionality where the number of features vastly exceeds the number of samples, it is likely to discover multiple sets of gene features that can discriminate samples from different groups of interests. To assess these candidate biomarker solutions, most biomarker discovery research like aforementioned studies have applied a single criterion, e.g., fold change in differential expression analysis or classification performance in cross-validation tests. In our previous study, we similarly highlighted a minimal biomarker panel that achieved the highest performance to discriminate different cellular states for *B. subtilis*. Individual scoring criteria, however, will always capture limited information and hence are inherently biased. There are multiple properties that a good biomarker panel for an in-vivo validation should present. Naturally good discriminative capacity between cellular states, but also biomarkers that are important to the gene regulatory control of the organism, to mention only two possible criteria. Therefore, there is a need for better panel selection methods that can rank candidate biomarker panels integrating a variety of data driven and domain-based criteria, so as to select a robust and reproducible biomarker panel.

One source of information that can be used to assess biomarkers, but has been often neglected, are models of gene regulation such as gene regulatory networks (GRN). A gene regulatory network is a collection of molecular species and their interactions, which together govern gene expression levels of mRNA and control gene-product abundance. GRNs play an important role in all kinds of cellular processes, including cell cycle, metabolism, and signal transduction. A well-curated GRN has already been described for *B. subtilis* [27]. This GRN has been rigorously validated experimentally and includes thousands of genes and hundreds of regulatory factors. GRNs such as this can reveal the regulatory circuits that are modulated by the identified biomarker genes, or which can, in-turn, regulate biomarker gene expression. Overall, by mapping biomarkers in a GRN it is possible to assess the diversity of cellular processes that the biomarkers are involved in.

As our current knowledge of regulatory interactions within the cell is still limited, co-expression networks (CEN), which are inferred from the correlations of expression patterns across different conditions [28, 29] can also provide additional information about transcriptional relationships between genes and their products. By grouping genes that are highly interconnected we can identify modules of genes with similar functions and relate the genes of unknown function to well-studied genes [30]. CEN can, therefore, be used to study the functional diversity of biomarkers as a supplementary method to complement GRNs.

We hypothesised that to identify a robust biomarker panel, a multimodal approach combining prior known information about gene interaction and co-expression with computational data-driven methods was crucial. We designed a two-step approach to rank and select a robust biomarker panel comprising key genes. Firstly, we applied the *Bacillus subtilis* biomarker identification model (BIM), we developed previously, to obtain a pool of candidate biomarker panels with satisfactorily high performance in predicting the stress state of a sample. Secondly, for each biomarker panel we integrated complementary information from BIM, GRN and CEN, developing a recommendation system to identify the biomarker panel with a few key genes to sense stress conditions. We successfully validated the robustness of the recommended panel on nine external datasets covering 10 stress conditions. These findings suggest that our *in silico* biomarker recommendation system, integrating multi-source knowledge and data-driven techniques, can facilitate *in vitro* experiments to sense cellular states for monitoring stress conditions in *B. subtilis*.

## 2 Methods

### 2.1 Datasets and data processing

For discovering biomarker panels, we used a tiling array dataset [31] that assays the transcriptomes of *B. subtilis* strain BSB1 measured under diverse conditions, including alternative nutrient shifts, lifestyle changes and adaptation to various stimuli. The wide range of conditions can lead to distinct transcriptional states in bacteria. Some of these conditions can cause bacterial stress, enabling the identification of biomarker genes specific to different stress states. Although, more recently, RNA-seq technology has also generated many transcriptomics datasets containing different conditions respectively, the integration of these small datasets was challenging since they are experimentally more diverse and would introduce between-experiment noise when included. The unified tiling array dataset from Nicolas and co-workers (referred to as the Nicolas dataset) was preferable for exploring the transcriptional profiles across conditions as between-experiment noises were minimised by conducting all experiments following standard operating procedures and estimating the gene expression quantities using the same signal processing protocols. The experimental procedures and signal processing protocols are described in Supplementary Material SOM1-2 from [31].

Nicolas and co-workers computed an aggregate expression index for each of 5875 transcribed regions as the median log2 expression signal intensity of probes lying entirely within the corresponding region. These expression values were pre-processed with quantile normalisation that makes the data distribution across samples identical to reduce between-sample variations. The raw data in Gene Expression Omnibus (GSE27219) and pre-processed data was made available at http://genome.jouy.inra.fr/basysbio/bsubtranscriptome.

To enable the discovery of relevant cellular states using unsupervised machine learning, we further processed data using the processing steps as elaborately explained in [26]. These steps included: a) filtering genes that were invariant across conditions; b) removing genes and samples related to late-stage sporulation conditions, as sporulation produces a very strong transcriptional response across a large number of genes that would mask many other cellular states we are interested in capturing; c) normalising the expression quantities by subtracting the corresponding reference conditions within each experiment; to produce a processed condition-dependent gene expression data (Figure 1a) for downstream analysis. The data processed using the above steps, containing 2536 genes and 180 samples, can be downloaded from https://github.com/neverbehym/transcriptional-biomarkers-subtilis.

**Figure 1.**
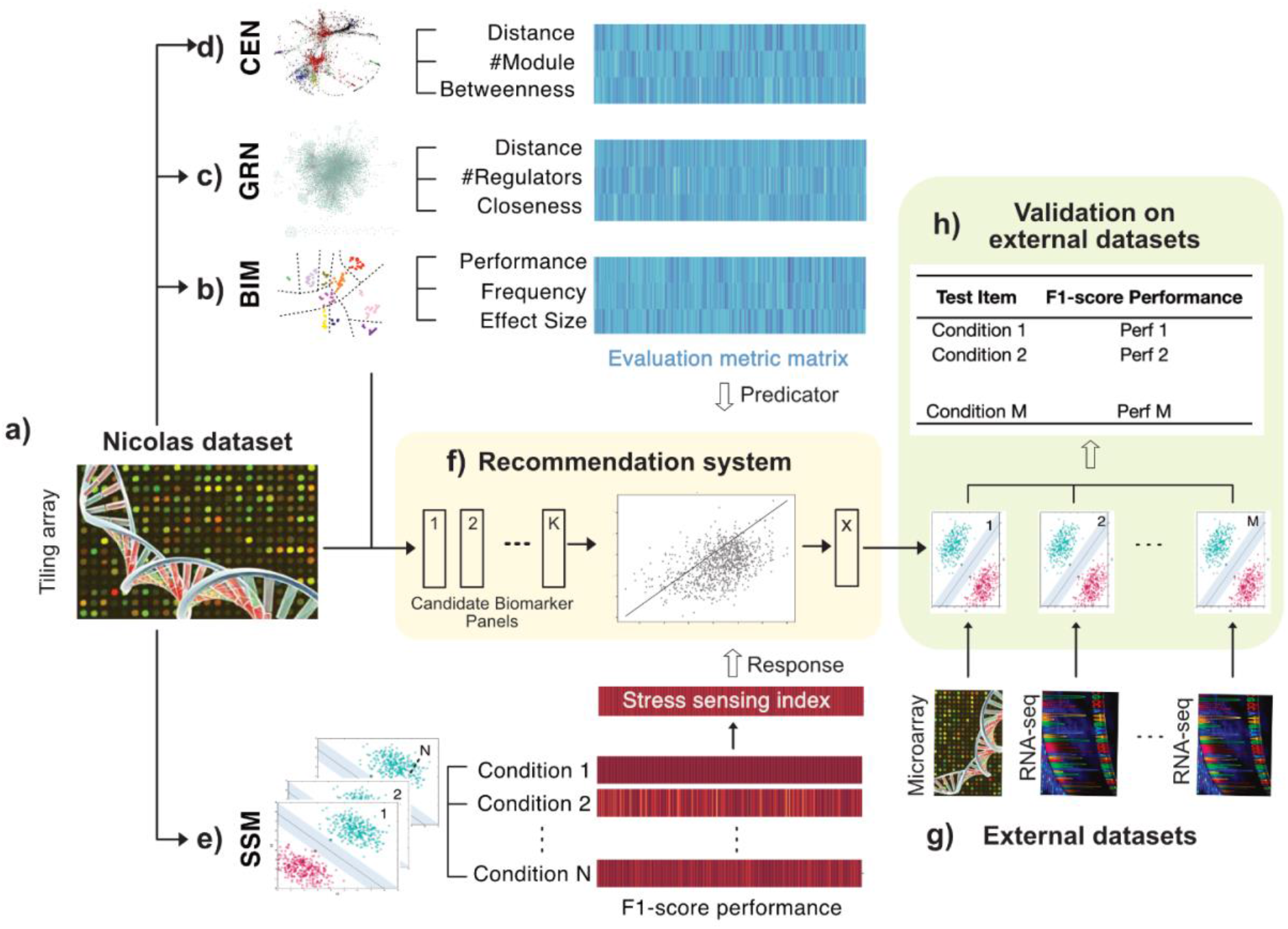
Overall approach. **a)** Nicolas dataset, which measures condition-dependent gene expression profiles using Tiling array technology, is used for biomarker identification and optimisation. **b) A** Biomarker Identification Model (BIM) is used to identify a pool of candidate biomarker panels and extract biomarker evaluation metrics: *Performance, Frequency, Effect Size*. **c) A** Gene Regulatory Network (GRN) is used to extract biomarker evaluation metrics: *Distance, #Regulator, Closeness*. **d) A** Co-Expression Network (CEN) is used to extract biomarker evaluation metrics: *Distance, #Module, Betweenness*. **e)** A Stress Sensing model (SSM) produces a stress sensing index for each candidate biomarker panel. The stress sensing index is calculated as the average f1-score performance on predicting N stress conditions in the Condition-dependent gene expression data. **f)** The recommendation system takes a set of candidate biomarker panels as input and recommends an optimal biomarker panel. This recommendation system is trained to relate evaluation metrics with the stress sensing index. **g)** A set of external datasets, collected in different centres with RNA-seq and Microarray technologies, are used for validation. **h)** Validation on external datasets is performed by assessing the performance of the recommended biomarker panel predicting M conditions included in the condition-specific gene expression datasets.

We also used nine condition-specific gene expression datasets (Figure 1g) for the validation of the biomarker identification model. These datasets, which are either accessible from Gene Expression Omnibus (GEO) or on request, were generated over past years using RNA-seq data or Microarray data in multi-centres including our own laboratory. Each of the datasets usually consists of mRNA samples collected under a test condition treated with a specific environmental perturbation as opposed to a control condition. The details of experimental conditions and GEO session IDs for all datasets used in this study can be found in Supplementary Table 1.

### 2.2 Overall approach

We proposed a in silico pipeline (Figure 1) to select a small and robust biomarker panel to sense a variety of stress states for *B. subtilis* We used the Nicolas dataset (Figure 1a) for training a recommendation system and a set of external datasets (Figure 1g) for validation of the recommended biomarker panel. Nine evaluation metrics were derived from a Biomarker Identification Model (Figure 1b), a Gene Regulatory Network (Figure 1c) and a Co-Expression Network (Figure 1d), and a stress sensing index was calculated by applying a Stress Sensing Model (Figure 1e). The recommendation system (Figure 1f) can select a biomarker panel from a pool of candidate panels by ranking the panels based on the evaluation metrics and coupling these evaluation metrics with the stress sensing index. We reported the performance of the selected biomarker panel to predict an extended list of conditions from external datasets for validation (Figure 1h).

### 2.3 Biomarker identification model

In our previous study [26], we proposed a Biomarker Identification Model (BIM), which was applied to the Nicolas dataset to discover biomarker panels indicative of the cellular states for *B. subtilis*. This BIM first discovers different cellular states introduced by a wide range of conditions using UMAP dimension reduction [32] and Leiden clustering [33] methods. The BIM then identifies panels of biomarkers indicative of these cellular states using our RGIFE [34], a recursive feature elimination-style feature selection algorithm. Here we improved the BIM to enable the identification of a sufficient number of biomarker panels of small size and high predictive performance by performing an additional feature elimination and evaluation process (Algorithm 1). On each of 500 biomarker panels of different sizes and prediction performances we identified from the previous paper, we iteratively removed individual genes while optimising cross-validation performance until reaching the desired panel size ranging from min feature size = 5 to max feature size = 10. Here the cross-validation performance was computed as the micro f1-score in 10-folds stratified cross-validation test. We then filtered the panels with cross-validation performance of less than perrformance_threhold = 0.9. By which we produced 949 candidate biomarker panels for downstream analysis.

#### Algorithm 1

Generation of candidate biomarker panels

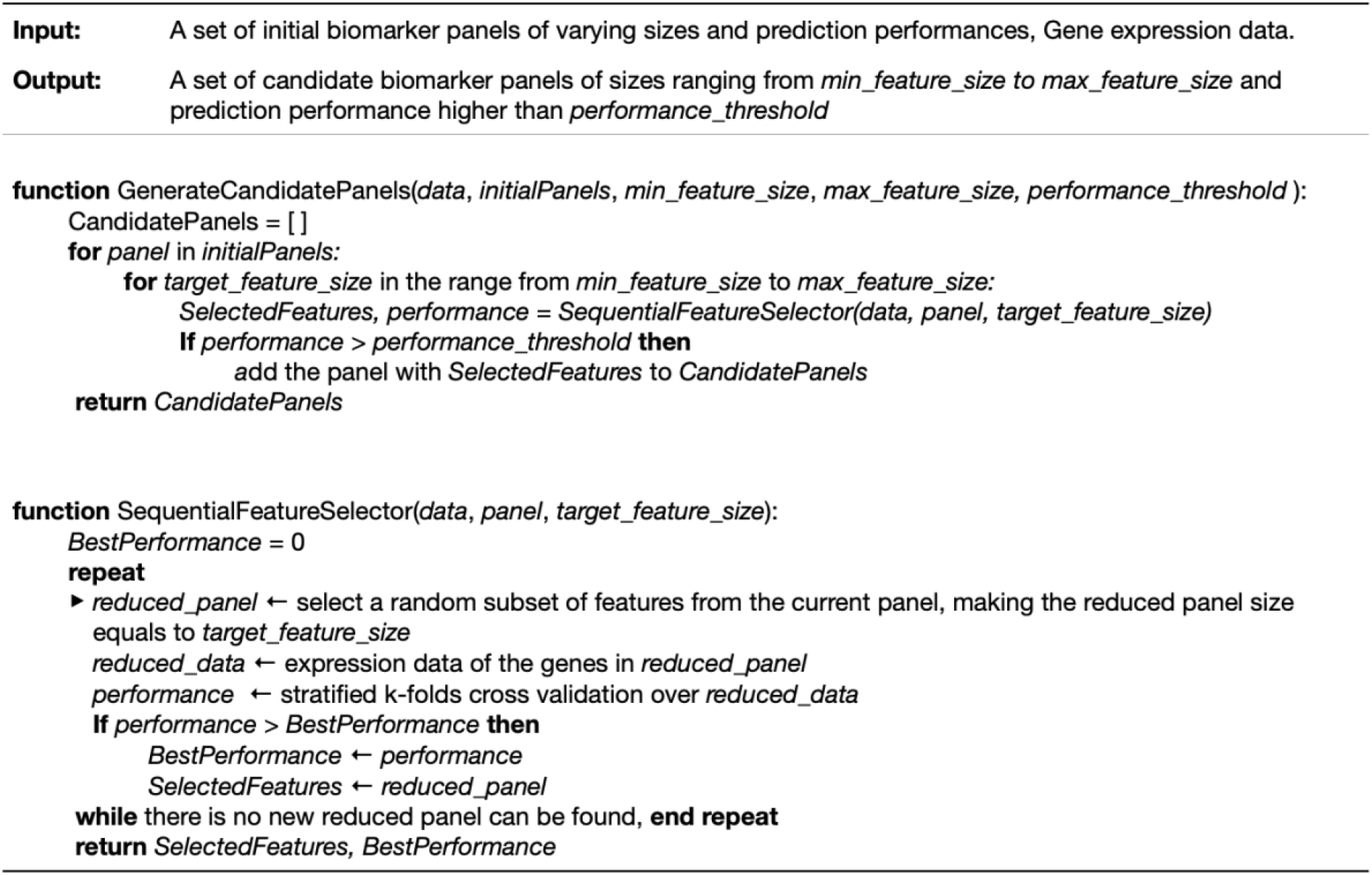

We extracted three metrics from this refined BIM to evaluate candidate biomarker panels (Figure 1b): a) *Performance* is used to measure the prediction performance of a given biomarker panel on distinct cellular states in the Biomarker Identification Model. It is calculated as 10-folds cross-validation f1-score for predicting ten cellular states with Random Forest classifier. b) *Frequency* reflects the consistency of a biomarker panel being selected across multiple repetitions of running BIM starting from different random states. As a high-throughput dataset tends to give false positive results, a biomarker panel with a higher *Frequency* value is expected to have less chance of being spurious signals. We first computed, for each gene, the frequency of appearing in the pool of candidate biomarker panels and then calculated Frequency as the average frequency values across all genes in a panel. c) *Effect Size* evaluates the overall strength of differential expressions in the biomarkers between distinct cellular states. A biomarker panel with a higher *Effect Size* value will likely make a reporter system with stronger signals. It is calculated as the average value of EffectSize_gene_ for all genes in each biomarker panel. As defined in Equation 1, 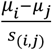 is used to compute Cohen’s d score, which measures the standardised difference in biomarker expression between two cellular states i and j.

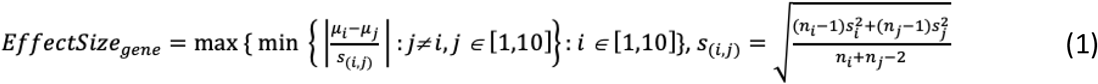

### 2.4 Gene Regulatory Network

To assess the diversity of cellular processes in which the biomarkers are involved with prior knowledge, we curated a gene regulatory network for *B. subtilis* by incorporating the network reconstructed by Jose and co-workers with the latest Subtiwiki regulon database. This network consists of 6704 edges, indicating the interactions between 294 regulators and 2816 targeted genes under various mechanisms such as transcriptional factors, RNA switches, riboswitches, and small regulatory RNAs (Supplementary Table 2). The network comprises 35 components, with the largest one containing 2798 nodes.

We extracted three evaluation metrics from GRN (Figure 1c): a) Number of regulators is the number of transcriptional regulators in GRN associated with the genes from a given biomarker panel. b) Distance indicates how distant any two genes from a given biomarker panel are in GRN. The distance between connected genes is calculated as the number of least hops in the network. The distance between unconnected genes is set as m+1, where m is the largest distance between any two connected genes in the network. c) Closeness is calculated as the average node closeness centrality across genes from a given biomarker panel in GRN.

### 2.5 Co-Expression Network

To study the functional diversity of biomarkers diagnostic of the transcriptional relationships discovered in GRNs, we constructed the co-expression network (CEN) that captures the expression similarity patterns across conditions. We applied weighted gene co-expression network analysis using the R package WGCNA [29].The resulting network is a fully connected and weighted network, with nodes being studied genes and edges reflecting the similarity of gene expression profiles across various conditions. First, we computed the adjacency value *a*_*(i,j)*_ as in Equation 2, where *s*_*(i,j)*_ is similarity strength based on Pearson correlation and β is soft-thresholding power index. We tuned the parameter β = 4 to maximise the scale-free topology criterion [35]. Second, we identified modules, i.e., clusters of highly connected genes, with the Dynamic Tree Cut method [36]. We tuned the parameters *detectCutHeight, mergeCutHeight, minModuleSize* in *blockwiseModules* function to maximise the average Overlap score across all modules in a network. The Overlap score for a given module is calculated in Equation 3, where *Overlap*_*m*_ measures the overlap level between *module*_*m*_ and a most concordant regulon in the GRN. We identified 55 modules with similar co-expression patterns as indicated by different colours, leaving 245 genes unassigned (grey). We summarised the profiles of these modules by studying the central genes with high intramodular connectivity, the highly overlapped regulons, and the overrepresented biological process in Gene Ontology. Please find the details in Supplementary Table 3.

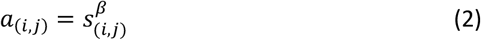

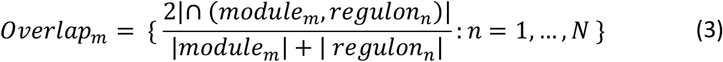

We extracted three evaluation metrics from the CEN (Figure 1d): The number of module (#Module) is the number of different modules in the CEN that are associated with the genes from a given biomarker panel. Distance measures the average distance between pairs of biomarker genes in the CEN. The cost of each edge is calculated as the inverse of co-expression strength, and the distance between two genes is set as the least sum of costs for the path connecting them. Betweenness is the average node betweenness centrality of biomarker genes in the CEN.

### 2.6 Stress sensing model

The Stress Sensing Model (SSM) is used to measure the overall performance of a biomarker panel in predicting stress states induced by a variety of test conditions (Figure 1e). While BIM identifies biomarkers indicative of different cellular states revealed by applying unsupervised machine learning methods on condition-dependent transcriptomes, SSM assesses the biomarkers using the condition labels provided. We first applied the Linear Support Vector Machine (SVM) model to classify gene expression patterns of a biomarker under each test condition from their gene expression patterns under the corresponding control condition. We computed the cross-validation f1-score performance to reflect the single-stress sensing power of this biomarker panel. We then computed the stress sensing index that reflects the multi-stress sensing power as the average f1-score performance across all test conditions. We applied SMM to measure the multi-stress sensing power of a pool of candidate biomarker panels identified by BIM, assessing the overall prediction performance in 13 test conditions included in Nicolas dataset (Table 1a).

**Table 1.** **a)** Conditions studied in the Nicolas datasets during the training process. **b)** Conditions studied in the external datasets during the validation process, in which some are similar to the conditions covered in the training dataset while some are different.

| a) | Test Conditions | Sample Size |  | b) | Test Conditions | Covered in Nicolas dataset | Sample Size |  |
| --- | --- | --- | --- | --- | --- | --- | --- | --- |
|  |  | Test | Control |  |  |  | Test | Control |
| Nicolas dataset | Anaerobic | 6 | 3 | External datasets | Antibiotic (amphotericin, SynAnt49, YydF) | N | 9 | 9 |
|  | Antibiotic (Mitomycin) | 6 | 6 |  | Cold | Y | 5 | 4 |
|  | Biofilm | 4 | 8 |  | Deep Starvation | N | 3 | 4 |
|  | Cold | 6 | 6 |  | Glycine betaine | N | 6 | 6 |
|  | Germination | 6 | 8 |  | Hydroxyurea | N | 6 | 6 |
|  | Heat | 6 | 6 |  | Heat | Y | 12 | 9 |
|  | Low motility | 6 | 8 |  | Oxidative (H<sub>2</sub>O<sub>2</sub>) | Y | 5 | 5 |
|  | Oxidative (Diamide, H<sub>2</sub>O<sub>2</sub>, Paraquat) | 15 | 6 |  | Potassium | N | 4 | 4 |
|  | Salinity | 6 | 6 |  | Pressure | N | 11 | 5 |
|  | Shift from glucose | 5 | 9 |  | Salinity | Y | 6 | 6 |
|  | Shift to glucose | 5 | 5 |  |  |  |  |  |
|  | Starvation | 9 | 8 |  |  |  |  |  |
|  | Stationary | 9 | 9 |  |  |  |  |  |

To mitigate the over-optimistic estimation of prediction performance due to limited dataset sizes, we performed 100 repetitions of the leave one out cross-validation and applied SMOTE [37] to oversample the minority class to the same size as the majority class in the training process.

### 2.7 Recommendation system

We built a recommendation system (Figure 1f) that takes a pool of candidate biomarker panels as input and produces a recommended panel as the output. We trained a regression model with Elastic Net regularisation (Equation 4, where *α* is the mixing parameter between L1 regularisation and L2 regularisation, *λ* is coefficient shrinkage parameter) predict the stress sensing index of a panel from 9 evaluation metrics described in sections 2.2 to 2.4. We tuned the model parameters *λ* = 0.2, *α* = 1e-4 to optimise the 10-folds cross-validation performance. A recommendation score was calculated as the predicted stress sensing index value (Equation 5) for each candidate biomarker panel. The recommendation system then selects the panel with the highest recommendation scores as the optimal biomarker panel.

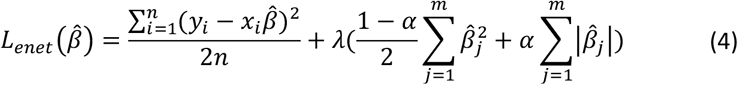

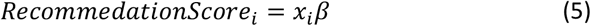

To understand the respective impact of each evaluation metric on the recommendation system, we calculated the SHAP values [38], reflecting the overall importance of each feature in a linear regression model. While model coefficients can also describe how the response values will change on the value of an input feature, the coefficients depend on the scale of the input features. Taking the distribution of feature values in regard, SHAP values are computed as the difference between the expected model output and the partial dependence plot at the feature’s value, and thus can better measure feature impacts on the model.

### 2.8 External validation

We ran external validation (Figure 1h) to test the robustness of the optimal biomarker panel selected by the recommendation system (see in 2.7) on several *B. subtilis* gene expression datasets independent of Nicolas dataset used for model training. We estimated the stress sensing power of the recommended biomarker panel specific to the conditions covered in these external datasets (Table 1b) by applying the stress sensing model (see in 2.6).

To assess the ability of the recommendation system to prioritise an optimal biomarker panel over other candidate biomarkers and random genes, we computed the stress sensing index for the remaining candidate biomarker panels and a set of random gene panels in comparison with the recommended biomarker panel. This set of random gene panels was generated by repeatedly random sampling an equal number of genes as in each candidate biomarker panel.

## 3 Results

### 3.1 Assessment of the evaluate metrics for candidate biomarker panels

We discovered 949 candidate biomarker panels that can discriminate 10 cellular states in *B. subtilis* by rigorously running the Biomarker identification model (see 2.3). These candidate biomarker panels have all achieved classification performance of more than 0.9 (f1-score) in cross-validation tests. The sizes of these panels, i.e., the number of genes each biomarker panel consists of, ranged from 6 to 10.

We derived a set of metrics that evaluated different properties of the candidate biomarker panels: *Performance* for classification performance to distinguish different cellular states; *Frequency* for the stability of genes being selected as a biomarker; *Effects Size* for the strength of distinguishability; *Number of regulators* (*#Regulators*) and *Number of modules* (*#modules*) for the diverse biological modalities in GRN and CEN; *Distance-GRN* and *Distance-CEN* for the coverage diameter in GRN and CEN; *Closeness* and *Betweenness* for the significance by centrality measurements in GRN and CEN. These evaluation metrics varied across candidate biomarker panels, with large dynamic ranges seen in the distribution (Figure 2a). The correlation analysis between these metrics and stress sensing index (Figure 2b) and the correlation analysis within these metrics (Figure 2c) indicated they are complementary measurements and that individually they are not sufficiently predictive of stress sensing power. The initial assessment of this set of selected metrics showed the potential in predicting the stress sensing power of the biomarkers by incorporating them in a computational model.

**Figure 2.**
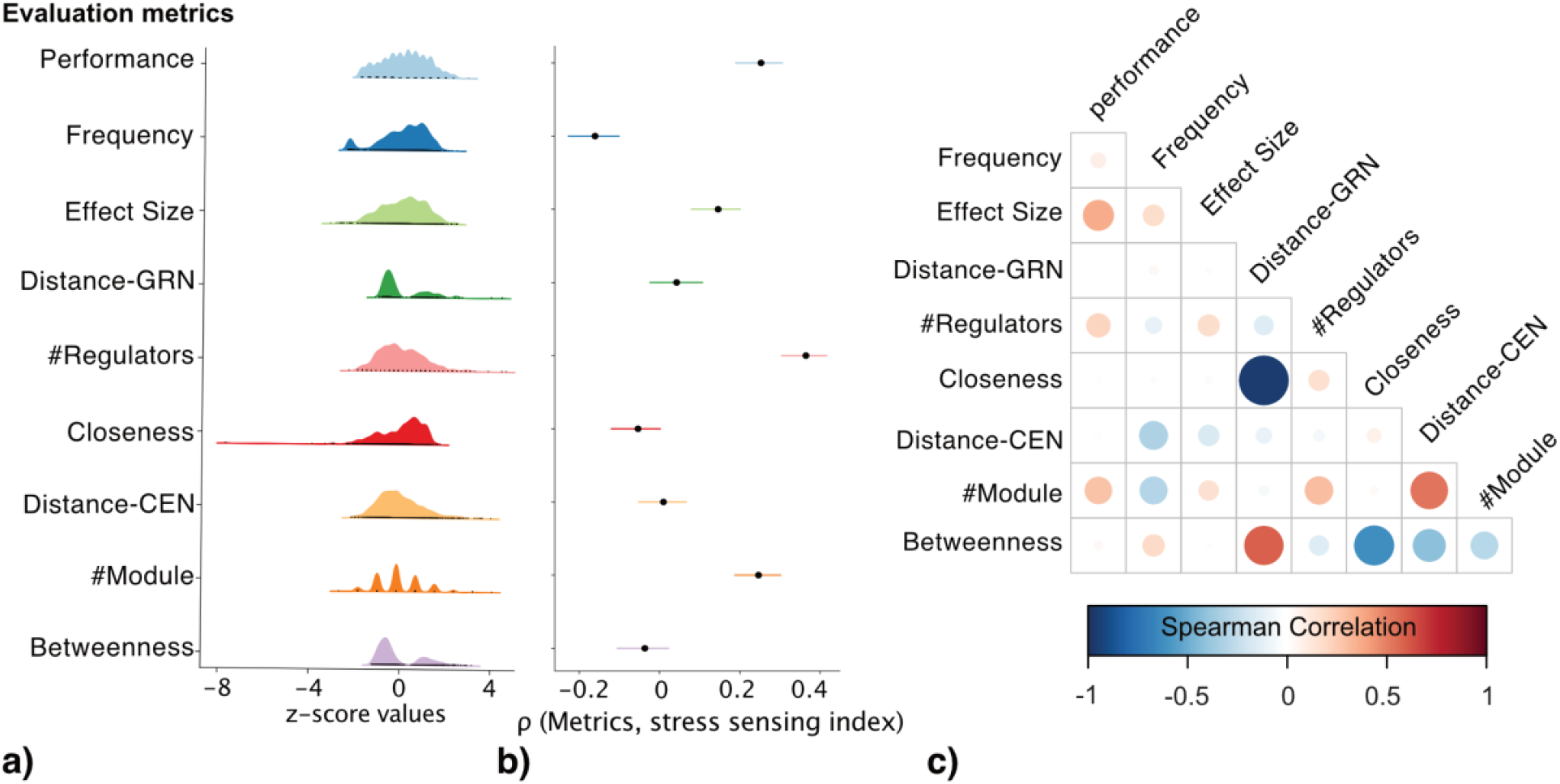
Evaluation metrics for biomarker panels. **a)** The distribution plots for different evaluation metrics. **b)** The Spearman correlation between each evaluation metric and stress sensing index is shown as the dot with the line indicates a 95% confidence interval. **c)** The correlation map between 9 evaluation metrics.

### 3.2 Recommendation system to select a robust biomarker panel

To build a recommendation system capable of selecting the optimal biomarker panel based on evaluation metrics and stress sensing index, we trained a regression model with Elastic Net regularisation to couple the evaluation metrics with the stress sensing index (Figure 3a-b). We observed a significant correlation (Spearman rho=0.44, p-value<0.001) between the actual stress sensing index and predicted stress sensing index produced by the trained regression model (Figure 3c). The trained model assigned a recommendation score as the predicted stress sensing index (Equation 5) to each candidate biomarker panel. Panel 661, which achieved the highest score, was selected as the recommended panel (Figure 3d). This recommended biomarker panel showed great prediction performance in classifying 13 stress conditions covered in Nicolas dataset, with mean f1-score achieved at 0.99 (i.e. acutal stress sensing index).

**Figure 3.**
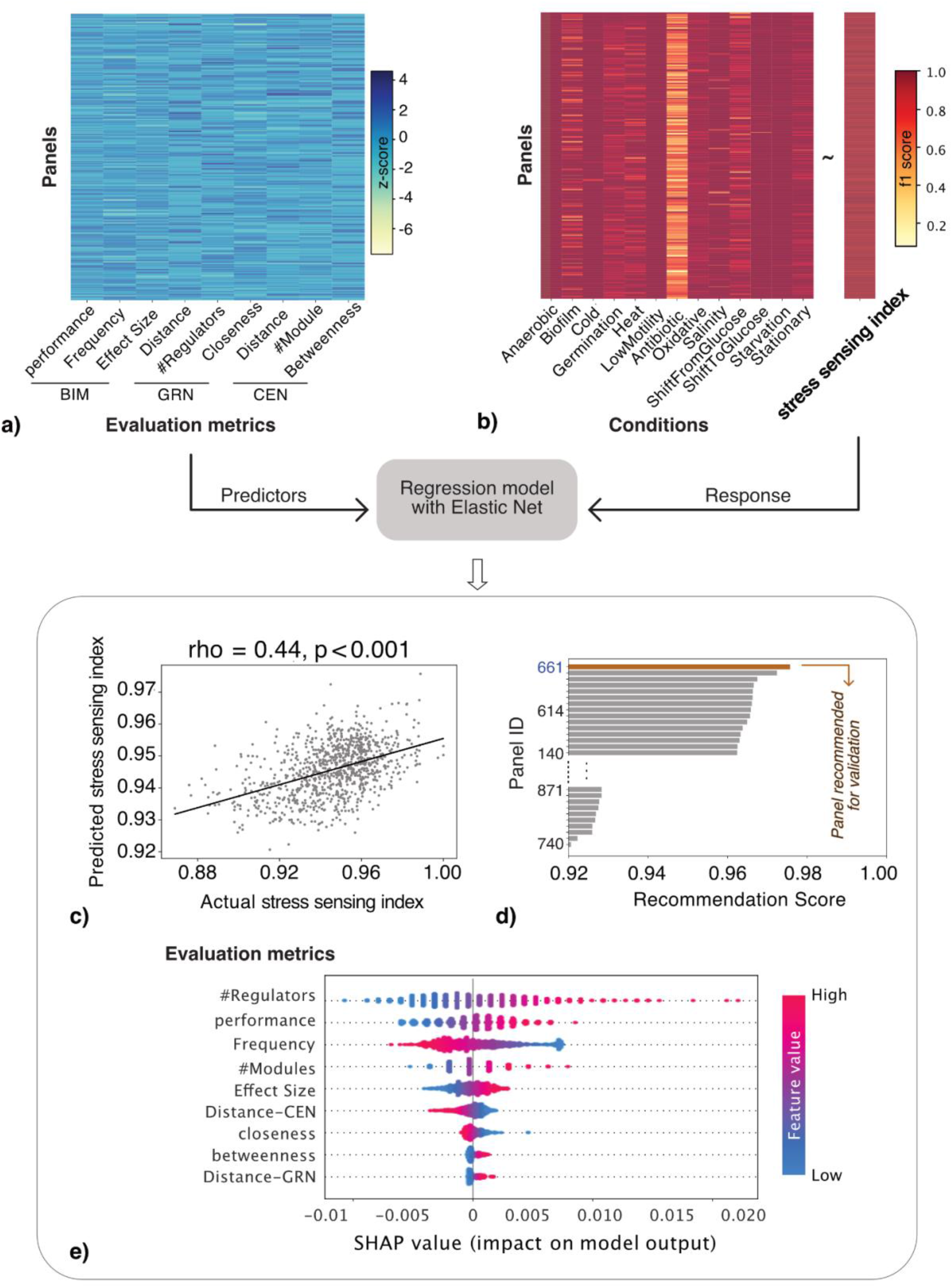
Recommendation of an optimal biomarker panel. **a)** The heatmap of 9 evaluation metrics for 949 panels. These metrics, derived from the Biomarker Identification model (BIM), Gene Regulatory Network (GRN) and Co-Expression Network (GEN), are Z-score standardised. **b)** The heatmap of classification performances (f1-score) in distinguishing 13 stress conditions respectively from control conditions, based on which stress sensing index is computed. The feature metrics of the biomarker panels are fitted in a linear regression model with Elastic Net regularisation to predict the stress sensing index based on 13 conditions. **c)** The scatter plot of predicted stress sensing index against actual stress sensing index. The Spearman correlation is 0.44 with significant effect (p-value < 0.001). **d)** The bar plot of *RecommendScore* in 949 candidate biomarker panels, sorted in descending order. The system selects Panel 661 with highest Recommend Score as the recommended biomarker panel. **e)** The SHAP summary plot of feature importance given by the trained regression model. All the instances are displayed as dots, with the colour indicating the original feature value and the x-axis SHAP value indicating the impact of this feature metric on the model.

We analysed the respective impact of each evaluation metric on the outcome of the recommendation system, i.e., recommendation score (Figure 3e). *Number of regulators*, which reflects how diverse the transcriptional regulatory parts are associated with the biomarker genes, showed the most significant impact on the model. The recommendation system gives a higher score for a biomarker panel consisting of genes that are involved in more regulatory parts. *Number of Modules*, which indicates the coverage of different modules in the co-expression network, similarly presented a positive correlation with the recommendation scores. *Performance, Frequency, Effect Size*, as the evaluation metrics extracted from the Biomarker identification model, respectively indicates the prediction power for classifying distinct cellular states, the uniqueness in candidate solutions, the strength of differential expression, were the remaining features among the top 5 contributors for ranking the candidate biomarker panels.

### 3.3 Validation of the recommended biomarker panel on external datasets

We validated the performance of the recommended biomarker panel on external datasets to classify samples grown under 10 test conditions from the samples grown under the corresponding control conditions (Table 1b). Despite variations observed in distributions of classification performance across candidate biomarker panels for all conditions, the recommended biomarker panel achieved 100% prediction accuracy for 7 conditions and still good prediction performance for the remaining 3 conditions (Figure 4a). Note that half of the conditions (e.g., Pressure, Hydroxyurea, Potassium, Glycine betaine) we tested here were not covered in Nicolas dataset used for the model training. This shows that the proposed knowledge-based recommendation system holds the potential of prioritising biomarker genes responsive to an extended set of treatment conditions that were not seen in the biomarker identification process.

**Figure 4.**
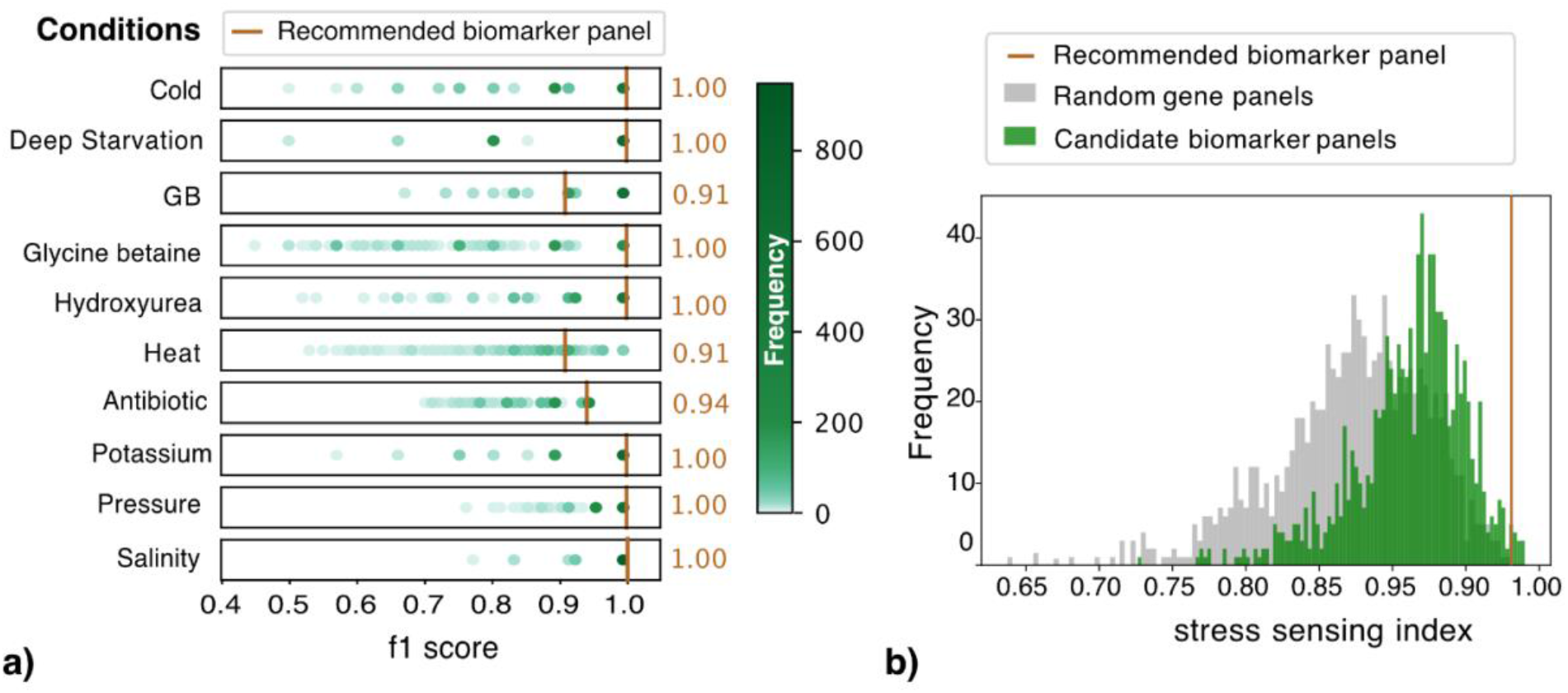
Validation performance of recommended biomarker panel on external datasets. **a)** A group of scatter histograms with each shows the distribution of validation performance (f1-score calculated on an external dataset) for a specific condition across 949 candidate biomarker panels. The hue of a dot reflects the frequency of occurrence for the corresponding value range. The validation performance of the recommended biomarker panel is marked as brown vertical lines with the value listed at the right for each condition. **b)** The grey histogram shows the distribution of stress sensing index across random gene panels, computed using external datasets. The green histogram shows the distribution of stress sensing index across candidate biomarker panels. A brown line marks the stress sensing index for the recommended biomarker panel.

**Figure 5.**
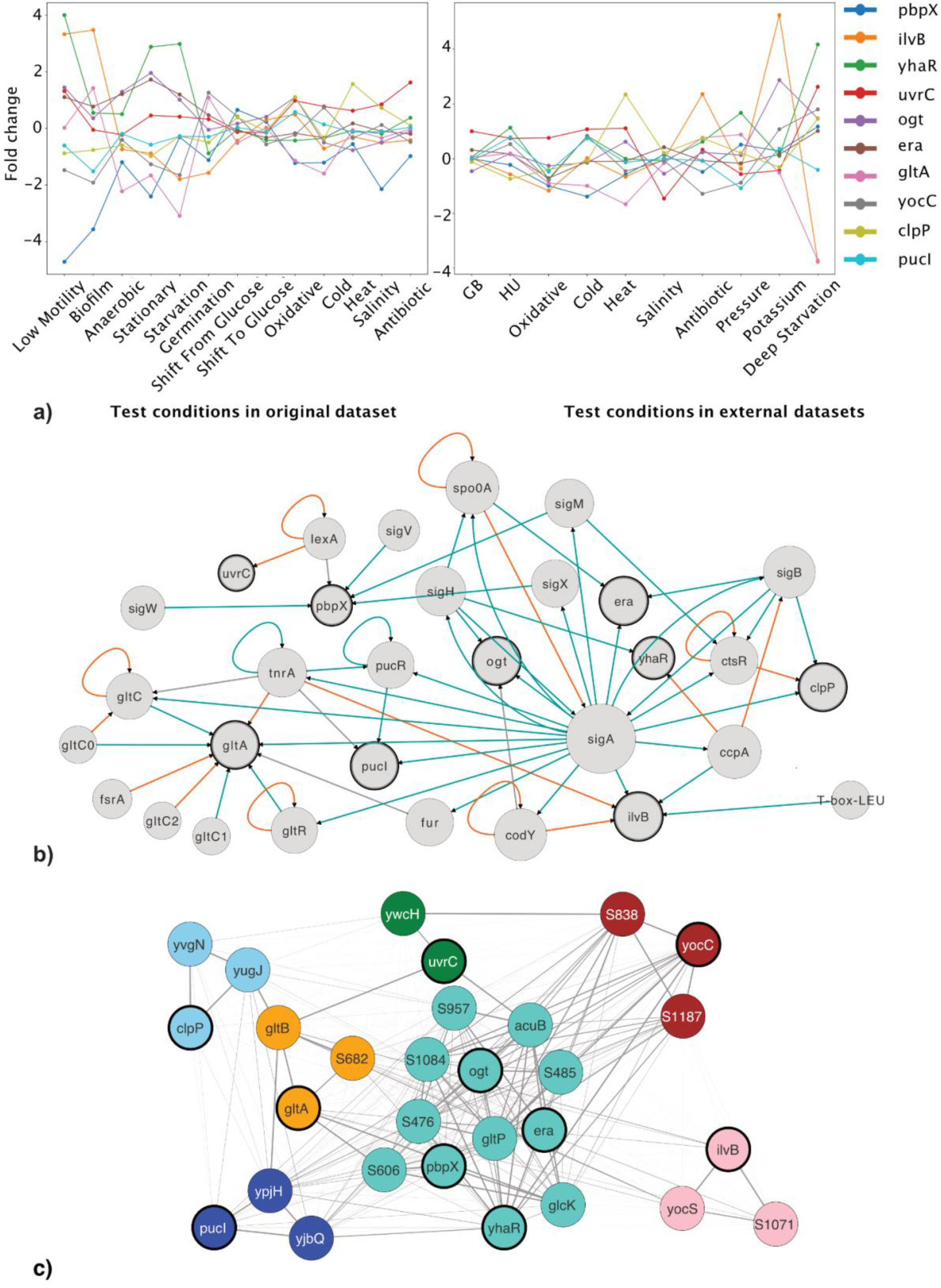
The functional profiles of recommended biomarker genes. **a)** The line charts show the fold changes of 10 biomarker gene expressions in test conditions against the control conditions. We tested 13 treatment conditions in the original training dataset and 10 in external datasets. **b)** The smallest component of Gene Regulatory Network that connects the recommended biomarker genes (highlighted in bold borderline). The node size is proportional to node degree centrality in this network. An edge indicates the regulation from a source node to a target node, with the orange line denoting suppression and green denoting activation. **c)** A subnet of Co-expression Network composed of recommended biomarker genes highlighted in bold borderline and their closest neighbour genes. The edge line width represents the adjacent value, thicker edge meaning higher similarity in co-expression patterns and longer distance in the network. The node colour indicates the specific modules to which it belongs.

To summarise the overall stress sensing power of a given biomarker panel, we averaged the prediction performance over all test conditions as the stress sensing index. We found that the stress sensing index for the recommended panel (0.98) is higher than 98% of the candidate biomarker panels and 99.5% of the random gene panels (Figure 4b).

### 3.4 Biological characterisation of the recommended biomarker panel

By training the recommendation system to predicate the stress sensing power of biomarker panels with evaluation metrics, we selected the biomarker panel with the highest predicted stress sensing index for in-vitro validation. As shown in Table 2, this recommended biomarker panel consists of 10 genes with various functions corresponding to DNA repair, endopeptidase activity, the synthesis of essential proteins, etc. These recommended biomarker genes presented varying alterations in transcription in response to different stress conditions, which combined can be used to distinguish a specific stress condition (Figure 4a).

**Table 2.** Gene functions and products of the recommended biomarker panel.

| Gene name | Gene function | Gene product |
| --- | --- | --- |
| yhaR | Unknown | putative dehydratase |
| ogt | DNA repair | O6-methylguanine DNA alkyltransferase |
| pbpX | endopeptidase | penicillin-binding protein X |
| gltA | glutamate biosynthesis | glutamate synthase (large subunit) |
| yocC |  | conserved hypothetical protein |
| era | ribosome assembly | GTP-binding protein |
| ilvB | biosynthesis of branched-chain amino acids | acetolactate synthase (large subunit) |
| uvrC | DNA repair after UV damage | excinuclease ABC (subunit C) |
| clpP | protein degradation | ATP-dependent Clp protease proteolytic subunit |
| pucl | purine utilization | allantoin permease |

The evaluation metrics *#Regulators* and *#Modules*, which have shown a significant impact on the stress sensing power of the biomarkers in the recommendation system, are relevant to the diversity of biological processes that biomarker genes are entailed. Therefore, studying involved regulatory parts and co-expression modalities can reveal the functional profiles of the biomarkers. We extracted the smallest component that connects the biomarkers from the Gene Regulatory Network, in which 20 sigma factors or transcriptional regulators were found associated with the regulations of these biomarker genes (Figure 4b). This includes *sigA* for household regulation, *sigB* for the general stress response, *sigH* for transiting to stationary growth phase, *sigM/sigV/sigX* for extracytoplasmic functions, *ccpA/codY/tnrA* for carbon, nitrogen or amino acid regulations, *lexA/ctsR* for DNA repair or heat shock response, etc. In complementary analysis, we also studied the Co-expression Network’s subnet consisting of biomarkers and their closest neighbours (Figure 4c). We found that 6 biomarker genes belong to unique modules while 4 are from the largest module (turquoise, 440 genes). As seen in the Supplementary Table 3B, these modules are associated with diverse biological processes, ranging from alternative carbon metabolism, various amino acid biosynthesis to protein repair and regulation of cell morphogenesis, etc.

## 4 Discussion

The objective of this study was to propose a computational method for selecting a robust biomarker panel that can be reproducibly applied in future studies to predict a wider range of stress conditions for *B. subtilis*. By quantitatively relating a collection of evaluation metrics that capture complementary information of the biomarker with the stress sensing power of the biomarkers in a regression model, we ranked the candidate biomarker panels based on the predicted stress sensing power and highlighted the evaluation criteria presenting the most impact to the recommendation system. The recommended biomarker panel showed prediction performance higher than 99.5% of the random biomarker panels and 98% of the candidate biomarker panels to predict 10 stress conditions in external datasets.

Although the accumulation of large-scale omics data recently has enabled the discovery of molecular biomarkers with high prediction performance, the high-dimensionality and high-noise characteristics of omics data have also posed challenges in stability and reproducibility of the machine learning methods applied. It is not uncommon to find little overlap between existing biomarker solutions produced by different datasets or different methods. The biomarker recommendation system we presented in this paper adopts a knowledge-based strategy that assesses the existing solutions using prior knowledge and multi-source criteria. The knowledge–based strategy and recommendation system proposed here can be readily applied to any biomarker discovery study where multiple biomarker solutions achieving comparable performance by traditional data-driven criteria, e.g., prediction accuracy, may be produced.

The key to successfully identifying biomarker solutions with improved robustness and confidence, however, lies in the selection of a set of effective evaluation metrics to assess biomarkers. In the case of identifying biomarkers indicative of various stress states in bacteria, it is important to utilise knowledge that can reveal the functional diversity of biomarker genes. Therefore, we incorporated the metrics extracted Gene Regulatory Network and Co-expression Network that are potentially relevant to diverse biological processes the biomarker genes may be involved in, which is likely the reason why the recommended biomarker panel showed great prediction accuracy in extended stress conditions including even conditions unseen in training data. We also want to note that this recommendation system can be adapted to incorporate any new evaluation metrics that are complementary and relevant.

While the recommended biomarker panel showed great prediction accuracy in diverse stress conditions from several independent datasets, the effectiveness of our results is constrained by the validation datasets used. Although we have maximally explored the relevant *B. subtilis* gene expression profiles that are publicly available for external validation, these datasets are all small, consisting of few samples grown under a single stress condition versus the corresponding control samples. They also vary on assay platforms (including RNA-seq and Microarray) and bacteria strains (ranging from wide type to BSB1 and JH642). Future studies are expected to validate these putative biomarkers in laboratory experiments with fine-controlled conditions.

To summarise, we showed that we could identify a robust biomarker that achieved improved prediction accuracy in external datasets by quantitatively evaluating the candidate biomarkers based on multi-source criteria incorporating various data-driven metrics and prior biological knowledge. Therefore, the in-silico methods we proposed in this paper can be applied to recommend an optimal biomarker panel indicative of various stress states in *Bacillus subtilis* for in vitro validation and implementation with increased confidence.

## Supporting Materials

Supplementary Table 1. Experiment Conditions in Training Dataset and Validation Datasets. Supplementary Table 2. Edge List of Gene Regulatory Network. Supplementary Table 3. Identified Modules in Co-expression Network.

## Author Contributions

Conceptualization, Y.H., J.B., A.W.; methodology, Y.H., N.S.; software, Y.H.; validation, Y.H., J.B., A.W., N.S.; formal analysis, Y.H.; data curation, investigation, and visualization, Y.H.; resources, Y.H., J.B., A.W.; original draft preparation, Y.H.; writing, review, and editing, Y.H., N.S, J.B., A.W.; supervision, J.B. and A.W.; project administration, J.B. and A.W.; funding acquisition, J.B. and A.W. All authors have read and agreed to the published version of the manuscript.

## Acknowledgements

The authors would like to thank Wendy Smith, David Markham and other ICOS members for valuable feedback and discussions. This research made use of the Rocket High Performance Computing service at Newcastle University.

